# ASER: Animal Sex Reversal database

**DOI:** 10.1101/2021.06.28.448629

**Authors:** Yangyang Li, Zonggui Chen, Hairong Liu, Qiming Li, Xing Lin, Shuhui Ji, Rui Li, Shaopeng Li, Weiliang Fan, Haiping Zhao, Zuoyan Zhu, Wei Hu, Yu Zhou, Daji Luo

## Abstract

Sex reversal, representing extraordinary sexual plasticity during the life cycle, not only triggers reproduction in animals but also affects reproductive and endocrine system-related diseases and cancers in humans. Sex reversal has been broadly reported in animals, however, an integrated resource hub of sex reversal information is still lacking. Here, we constructed a comprehensive database named ASER by integrating sex reversal-related data of 18 species from Teleostei to Mammals. We systematically collected 40,018 published papers and mined the Sex Reversal-associated Genes (SRGs), including their regulatory networks, from 1,611 core papers. We annotated homologous genes and computed conservation scores for whole genomes across the 18 species. Furthermore, we collected 206 available RNA-seq data and investigated the expression dynamics of SRGs during sex reversal or sex determination processes. In addition, we manually annotated 551 ISH images of SRGs from the literature and described their spatial expression in the gonads. Collectively, ASER provides a unique and integrated resource for researchers to query and reuse organized data to explore the mechanisms and applications of SRGs in animal breeding and human health. The ASER database is publicly available at http://aser.ihb.ac.cn/.

## Introduction

Sex determination mechanisms in animals mainly include genetic sex determination (GSD) and environmental sex determination (ESD) [1]. In GSD, the primary sex of organisms is determined by genetics during fertilization, while organisms with ESD remain bipotential gonads until they perceive environmental stress to promote sex differentiation during ontogeny [2]. For many years, it was dogma in vertebrates in the field of sex determination that sex would be fixed for life after primary sex determination. After sex reversal was first reported in *Aplocheilus latipes* and natural sex reversal was found in *Monopterus javanensis* [3], it has been widely accepted that sex determination is amazingly plastic in vertebrates, especially in fish. This plasticity shows that sexual fate is not an irreversible process. Indeed, this reversible process leads to sex reversal, a redirection of the sexual phenotype during development [4]. Environmental factors can override genetic factors to redirect sexual fate in fish [5] and reptiles [6]. Sex reversal was found to be driven by diverse factors, such as genes, hormones, temperature, and social changes [7]. Unlike sex change, which implies a transition from the stabilized sex to the opposite sex, sex reversal occurs during gonadal development, including the initiation phase and maintenance phase of sex determination [4].

Specifically, sex reversal has been studied in fish, reptiles, birds, amphibians and even in mammals. In fish, gonadal differentiation is roughly divided into two groups: hermaphroditic and gonochoristic [5]. Hermaphroditic species undergo sex reversal during their lifetime and include 3 strategies: female-to-male (protogynous), male-to-female (protandrous), or bidirectional (serial) sex change [8]. Taking *Monopterus albus* as an example, an individual is female from the embryonic stage to first sexual maturity, then enters an intersex state, and later develops into a male [9]. Additionally, some hermaphroditic species undergo socially cued female-to-male sex reversal, whereby the removal of the dominant male induces sex reversal in a resident female, such as *Thalassoma bifasciatum* [10]. Among gonochoristic fish, sex reversal is a synergistic result of both GSD and ESD [11]. For example, *Cynoglossus semilaevis* is a gonochoristic fish with a female heterogametic sex determination system (ZW♀/ZZ♂) characterized by GSD and TSD (a subclass of ESD) [12]. In many reptiles, including *Trachemys scripta*, gonadal sex is determined by the environmental temperature during egg incubation [13]. However, estrogens, including estradiol-17β, have also been proven to participate in the sex determination of *T. scripta* [14]. Sex reversal in birds such as *Gallus gallus*, is mainly related to alterations in sex steroid hormone action, especially estrogens [15]. Amphibians also show plasticity in sex determination, influenced by estrogens, androgens [16], and sometimes by temperature [17]. Sex determination in mammals has been reported to depend on three processes: chromosome determination (XX or XY), appropriate pathway of gonadal differentiation, and accurate development of secondary sexual characteristics [18]. Disrupting any of these three steps of gonadal differentiation can lead to aberrant sex determination. In *Homo sapiens*, the frequencies of XX and XY sex reversal are 1/20,000 and 1/3,000, respectively, and most of these cases are caused by translocations of the sex-determining *SRY* gene [19]. Although sex reversal has been broadly reported among vertebrates, the molecular events underlying sex reversal remain poorly understood, limited by the lack of integrated omics data across species.

Although there are several reproduction-related resources, such as GUDMAP [20], GonadSAGE [21] and ReproGenomics Viewer [22], an integrated and dedicated database for the community studying sex determination and differentiation is missing. The GUDMAP database is a comprehensive gene expression dataset of the developing genitourinary system in mouse with both *in situ* and microarray data. GonadSAGE is a serial analysis of gene expression database for male embryonic gonad development in mouse. The ReproGenomics Viewer is a cross-species database of omics data such as RNA-seq and ChIP-seq for tissues related to reproduction, such as gametogenesis, in 9 model organisms. Here, we developed the Animal Sex Reversal database (ASER), the first functional genomics hub for sex reversal to our best knowledge. The main works of ASER can be roughly divided as follows: (1) We screened 18 important and typical species with sex reversal phenomena from Teleostei to Mammalia, including *Betta splendens, Cyprinus carpio, Cynoglossus semilaevis, Danio rerio, Equus caballus, Epinephelus coioides*, *Lates calcarifer, Monopterus albus, Oryzias latipes, Oreochromis niloticus, Paralichthys olivaceus, Thalassoma bifasciatum, Xenopus laevis, Gallus gallus, Bos taurus, Mus musculus*, and *Homo sapiens*, and summarized the major inducements of sex reversal or common approaches used to manipulate sex in these species (**Table 1**). (2) We compiled a list of the most common genes or drugs related to sex reversal. Then, we collected and analyzed PubMed literature to mine the Sex Reversal-associated Genes (SRGs) and obtained their regulatory networks. Meanwhile, we gathered protein-protein interaction networks related to SRGs from the STRING database. (3) To facilitate users comparing the homology of SRGs in different species, we collected or assembled the gene annotations for the 18 species, identified homologous genes, computed the basewise conservation scores across these species, and identified conserved motifs for orthologous gene groups. (4) We systematically processed available RNA sequencing (RNA-seq) data and provided gene expression dynamics during sex reversal between females and males or different developmental stages. A user-friendly genome browser was customized to visualize these genome-wide data. (5) We collected and annotated available *in situ* hybridization and immunocytochemistry (ISH, FISH, and ICH) data to display the spatial expression of SRGS in the gonads. In conclusion, our ASER database provides comprehensive and systemic integration of sex reversal related data, and we believe that this open resource will greatly promote research on the mechanisms of sex reversal.

**Table 1.** Inducements of sex reversal or common approaches used to manipulate sex in 18 species.

| Species | Gonadal differentiation | Inducements/Approaches |
| --- | --- | --- |
| *Cyprinus carpio* | gonochoristic | H/D; TEM |
| *Danio rerio* | gonochoristic | GA; H/D; TEM |
| *Oryzias latipes* | gonochoristic | GA; H/D; TEM |
| *Oreochromis niloticus* | gonochoristic | GA; H/D; TEM |
| *Epinephelus coioides* | hermaphroditic | NAT; GA; H/D; SF |
| *Thalassoma bifasciatum* | hermaphroditic | NAT; SF |
| *Betta splendens* | gonochoristic | H/D |
| *Monopterus albus* | hermaphroditic | NAT |
| *Lates calcarifer* | hermaphroditic | NAT |
| *Paralichthys olivaceus* | gonochoristic | H/D; TEM |
| *Cynoglossus semilaevis* | gonochoristic | NAT; TEM |
| *Xenopus laevis* | gonochoristic | H/D |
| *Trachemys scripta* | gonochoristic | H/D; TEM; GA |
| *Gallus gallus* | gonochoristic | H/D |
| *Homo sapiens* | gonochoristic | GA |
| *Mus musculus* | gonochoristic | GA |
| *Bos taurus* | gonochoristic | GA |
| *Equus caballus* | gonochoristic | GA |

## Data collection and database content

### Framework of ASER

An overview of the ASER database and web server is shown in **Figure 1**. The ASER database contains information for 18 sex reversal species, SRG regulatory networks, homology alignment, and (fluorescence) *in situ* hybridization and immunocytochemistry (ISH, FISH, and ICH) images of SRGs. The preprocessed data was managed with the MySQL database. Django-based applications were developed to provide a user-friendly interface including an embedded genome browser for visualizing genome-wide data. The key workflows, tools, and processed data are summarized in Figure S1A-B and described in detail below.

**Figure 1.**
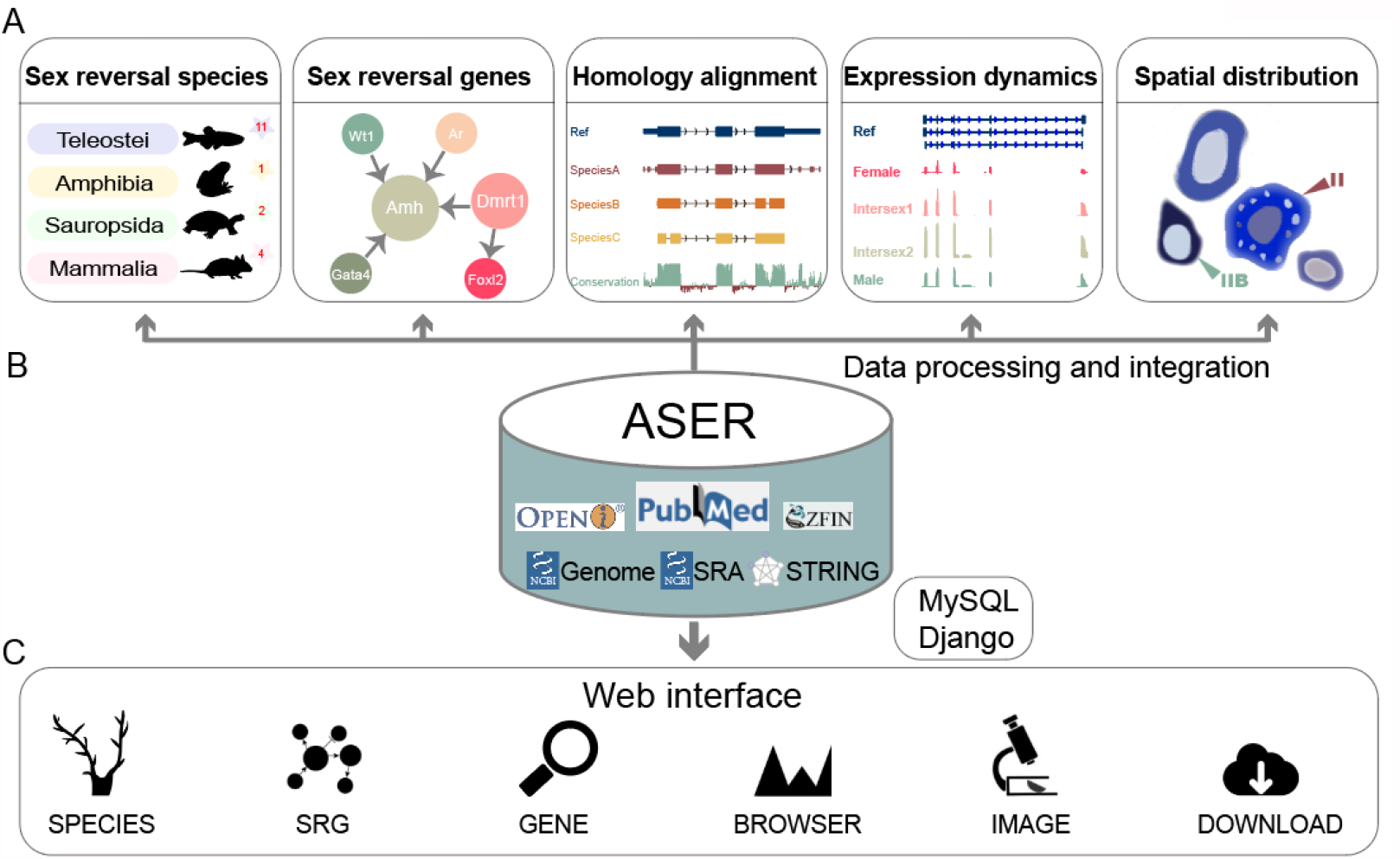
Schematic diagram of ASER database. **A**. Five main functional modules in ASER, including the 18 sex reversal species, sex reversal genes (SRGs) and their regulatory networks, multiple sequence alignments and conservation scores, gene expression dynamics during sex reversal from RNA-seq data, spatial distribution of SRGs from ISH, FISH and ICH images. **B**. Data sources in ASER database. ASER stores all processed data in a MySQL database with additional indexes and uses the Django framework for interactive queries from the web interface to the backend database. **C**. Overview of the ASER web interface. The main functionalities are provided and organized into six modules.

### Data sources

SRG information and their regulatory networks were curated from PubMed literature. All genome sequences and species information used in this database were downloaded from NCBI public databases. All raw sequencing data were downloaded from the Sequence Read Archive (SRA) of NCBI. The sets of RNA-seq data were organized by species, gonad developmental stages, and temperature (Table S1). In addition, we retrieved images related to sex reversal from the OPENi (https://openi.nlm.nih.gov/) and ZFIN databases [23].

### SRG mining

We retrieved thousands of articles from PubMed by querying species and functional keywords (e.g., sex reversal). First, the abstracts and full texts of these articles were collected by text crawler technology. The full texts of non-open access papers were obtained through the library portal of Wuhan University. Next, we separated a chunk of continuous text into separate words, and carried out word stemming to remove plural and different tenses. Then, we removed stop words such as such as “the”, “is”, and “however”. Finally, we counted the frequency of the words from the literature and manually filtered out some high-frequency but irrelevant words such as “masculinizing”, “ovotestes”, “pseudomale”, “hermaphrodite”, and “gynogenesis” into a blacklist until most of the high-frequency words were gene symbols and drug names. The remaining words related to genes and drugs were manually added to the wordlist (Table S2).

We retained 1,611 papers that contained the words in the wordlist and manually read them with notations about SRG regulation (Table S3). Finally, we found 258 SRGs, 6 drugs and 11 hormones, which were validated to be functional in sex reversal in different species, and constructed the regulatory networks of SRGs. We next predicted another 498 genes that were homologous with those SRGs in the 18 species (Figure S1C). Furthermore, protein-protein interaction (PPI) networks of SRGs were extracted from the STRING database [24].

### RNA-seq data processing

The data quality of the collected RNA-seq data was assessed using FastQC (http://www.bioinformatics.babraham.ac.uk/projects/fastqc/), and the adapters and low-quality bases in raw reads were removed using Trim Galore (http://www.bioinformatics.babraham.ac.uk/projects/trim_galore/). Filtered reads were aligned to the genome using STAR [25] in end-to-end mode. The primary alignments were retained through SAMtools [26]. Gene expression quantification in FPKM (Fragments Per Kb of exon per Million mapped fragments) was computed using StringTie [27]. Differential expression analysis was performed using DESeq2 [28].

### Transcriptome assembly

High-quality reads were *de novo* assembled using StringTie [27] with default parameter settings. The longest ORFs were predicted in assembled transcripts using TransDecoder.LongOrfs (https://help.rc.ufl.edu/doc/TransDecoder). DIAMOND [29] was used to collect homologous evidence of identified ORFs from the UniProt database (https://www.uniprot.org/). The potential coding regions were further refined by TransDecoder.Predict. Finally, a GFF3 file based on the coding regions of the reference genome was generated through the cdna_alignment_orf_to_genome_orf.pl function in TransDecoder.

### Homology alignment

Orthologous groups of SRGs were identified among all sex reversal species using the BLAST [30] all-v-all algorithm in OrthoFinder [31]. Conserved motifs of orthogroups were predicted using MEME [32]. Comparisons between conserved motifs and known motifs were performed using Tomtom [33]. Species tree was constructed according to the species taxonomy on NCBI. Meanwhile, the evolutionary relationship was verified by OrthoFinder using the STAG [34] algorithm and rooted using the STRIDE [35] algorithm.

For the orthologue tracks in a reference species, the homologous genes in other species were mapped to the reference genome using Blat [36]. Alignments with sequence identity larger than or equal to 60% were retained, and the maximum intron size was set to 450,000 bp.

For the conservation track, pairwise alignments between genome sequences were built using LASTZ [37], and MULTIZ [38] was then used to construct multiple alignments, based on which the conservation scores were calculated using phyloP from the PHAST package [39].

### Image collection and annotation

We collected available (fluorescence) *in situ* hybridization and immunocytochemistry (ISH, FISH and ICH) data related to SRGs from the OPENi and ZFIN databases. The images were classified by gene, differentiation status, developmental period, and gender. We manually added descriptions for those images based on the original figure legends and articles.

### Web interface and usage

ASER is a user-friendly database, and all the contents are interactive and dynamic. There are five main functional modules, including SPECIES, IMAGE, SRG, GENE and BROWSER. In addition, the SEARCH module was developed to display and interconnect different kinds of data in other modules.

For the “SPECIES” module, the evolutionary tree constructed for the 18 sex reversal species is displayed on the main page (**Figure 2A**). The reported inducements of sex reversal or common approaches used to manipulate sex in each species are displayed on this page, including natural processes, genetic abnormality (*e*.*g*., *amh* overexpression), administration of exogenous hormones or drugs (*e*.*g*., 17α-methyltestosterone), temperature changes during gonadal differentiation, and manipulation of social factors. The literature supporting this information is also provided. Users can click on any species to obtain detailed descriptions and genome information for this species (**Figure 2B**).

**Figure 2.**
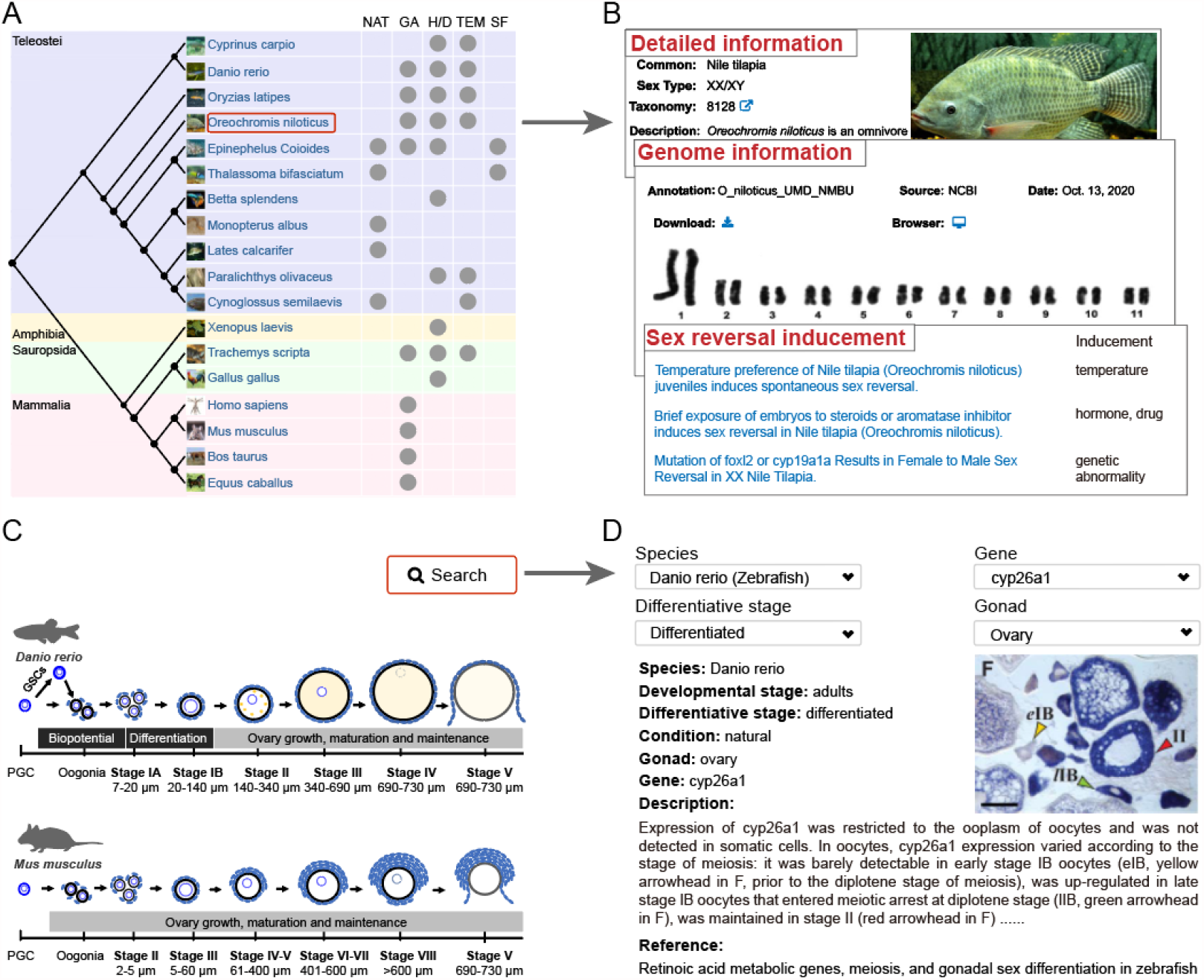
Species and image modules in ASER. **A**. Evolutionary relationship and sex reversal inducements of the 18 species belonging to Teleostei, Mammalia, Sauropsida, and Amphibia. NAT: Natural; GA: Genetic abnormality; H/D: Hormones or drugs; TEM: Temperature; SF: Social factors. **B**. Detailed description, genome information, and references related to sex reversal in each species. **C**. Examples of gonadal morphology at different developmental stages in mouse and zebrafish. **D**. Image page describing the spatial distribution of SRGs in gonads, with a representative example of cyp26a1 in ovaries.

For the “IMAGE” module, we summarized the morphological characteristics of zebrafish and mouse ovary at different developmental stages to help users better understand the content of this module (**Figure 2C**). The (fluorescence) *in situ* hybridization and immunocytochemistry (ISH, FISH and ICH) data related to specific SRGs can be queried in different ways by species, gene, differentiative stage, and gonad. Detailed descriptions of images are shown to help users understand the spatial distribution of SRGs in the gonads (**Figure 2D**).

The “SRG” module includes word cloud, regulatory and PPI networks, and search pages. The word cloud figure is dynamically presented by species with hyperlinks on the nodes (**Figure 3A**). When the user clicks one node, the original references and additional actions for more information will be shown under the figure. For any validated SRG, ASER allows users to obtain its regulators (including genes, hormones, drugs), targets, and the associated modes of regulation (**Figure 3B**). At the same time, PPI networks of these SRGs in different species are also displayed (**Figure 3C**), in which the colors of the edges are used to distinguish known interactions (experimentally_determined_interaction, database_annotated), predicted interactions (neighborhood_on_chromosome, gene_fusion, phylogenetic_cooccurrence), and other types (homology, coexpression, automated_textmining). In addition, the search page provides an interface for a specific SRG to show its regulatory network and more detailed information, such as tissue, developmental stage, and literature evidence (**Figure 3D–E**).

**Figure 3.**
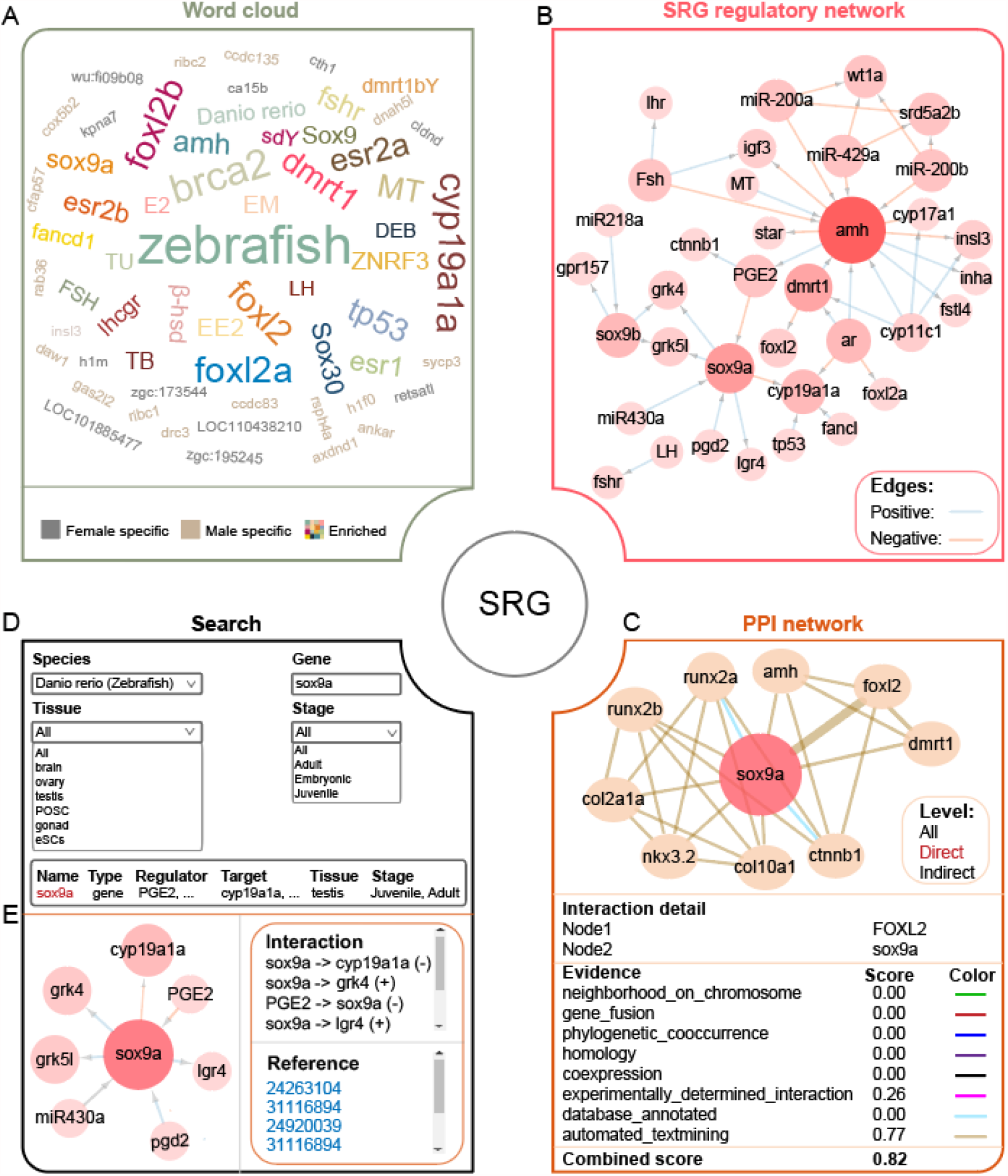
SRG module in ASER. **A**. Word cloud for genes or drugs mined from the literature as well as the top 50 genes that are specifically expressed in females or males. The size represents word frequency. **B**. Representative regulatory network of SRGs in *zebrafish*. The size represents the node degree in the network. **C**. An interactive view of the Protein-Protein Interaction (PPI) network for the SRG query. **D**. SRG regulatory network search page by species, tissue, gene and stage. **E**. Detailed information, regulatory relationships, and literature evidence for the SRG query.

The “GENE” module provides different kinds of data for any annotated gene, some of which are linked to the “BROWSER” module for visualization. The links corresponding to the query gene are shown in the search page by species and gene symbol (**Figure 4A**). Detailed information for a specific query gene includes its orthogroup in all species, predicted motifs, and similar known motifs (**Figure 4B**). The orthologous genes and 18-way conservation scores for the query gene can be inspected in BROWSER tracks (**Figure 4C**). Detailed alignment information can be obtained and downloaded by clicking on the track. In addition, the gene expression quantifications in FPKM across different stages, tissues, and conditions are shown as bar plots and in detail as tables (**Figure 4D**). The RNA-seq signal profiles are displayed in BROWSER, and the tracks can be customized easily, including color, scale, height, and *etc*. For any species, the available tracks can be dynamically selected or unselected. For example, in *Danio rerio*, a subset of RNA-seq tracks are shown for the *sox9* gene during sex reversal (**Figure 4E**).

**Figure 4.**
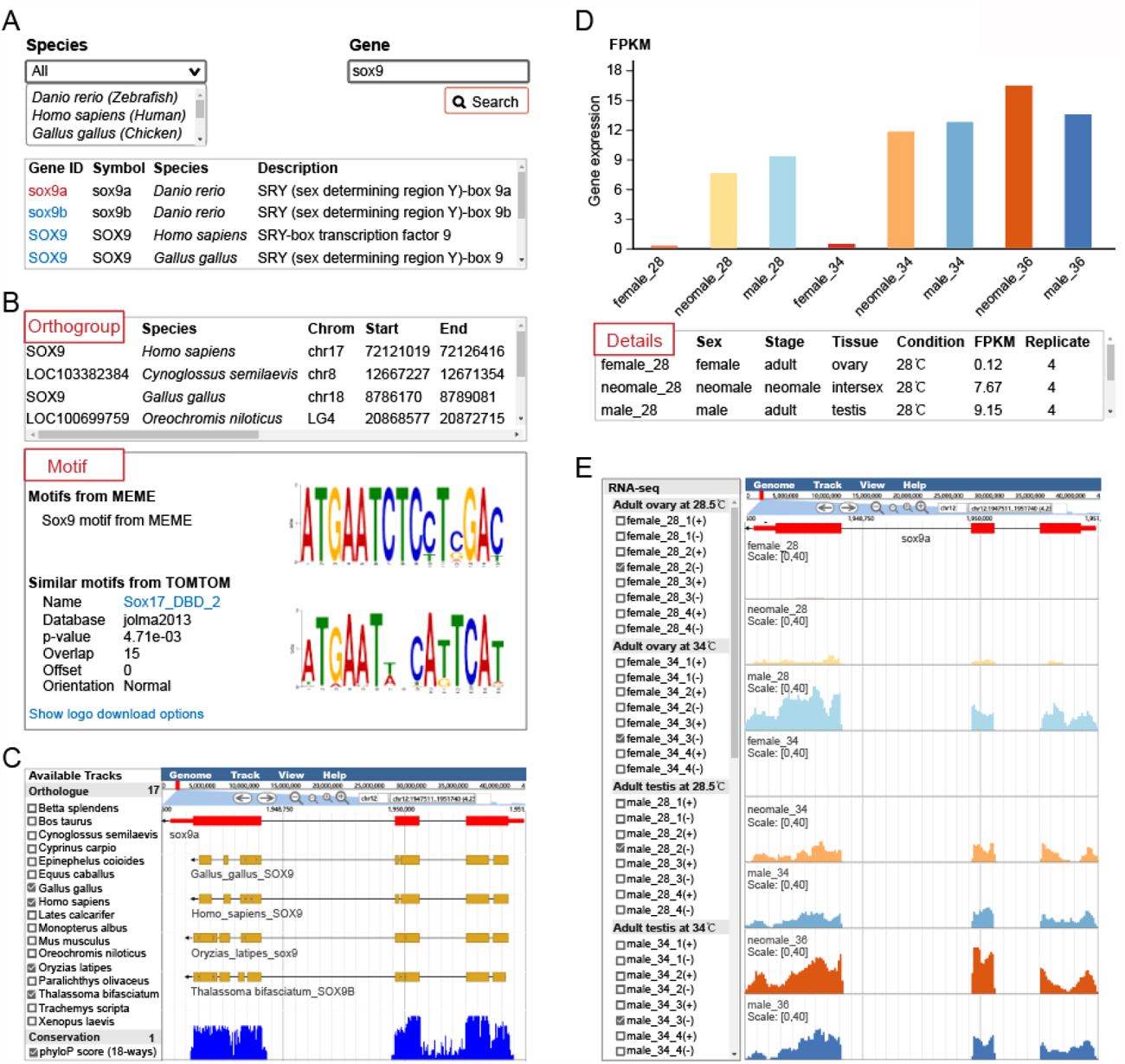
Gene conservation and expression modules in ASER. **A**. Search page for any gene in the 18 species. **B**. Homologous genes in the 18 species (top) and motifs corresponding to the query gene (bottom). The conserved motif for these homologous genes was computationally identified by MEME, and known motifs similar to this motif were also presented. **C**. Genome browser tracks of orthologue and conservation scores (18-ways). **D**. Expression dynamics of representative query gene. **E**. Genome browser view of processed RNA-seq signals for a representative query gene (*sox9a* in *zebrafish*).

## Discussion

Studies on sex reversal have been especially useful in helping redefine the concept of sex determination. There are diverse master sex-determining genes reported in different species. In addition, genes previously known to be involved in sex determination or differentiation are emerging as potential key components of sex reversal in other vertebrates [40]. Therefore, studies in different species continue to reveal genes with unexpected roles in sex reversal, and their homologs in other vertebrates also deserve investigation. ASER fills the gap of the sex reversal database by integrating diverse information at different levels for the 18 species with sex reversal, including curated SRGs, RNA-seq data, image data, and conservation data.

For any collected sex reversal species, users can obtain major inducements of sex reversal in this species in the SPECIES module. For any SRG, users can obtain its regulators, targets during sex determination in the SRG module and spatial distribution in different stages in the IMAGE module. For any annotated gene, users can obtain its homologous genes and conserved motifs in 18 species in the GENE module. Furthermore, users can also explore and visualize expression dynamics across different conditions in the GENE or BROWSER modules.

In the future, we will continuously select important and typical sex reversal species as their complete genome and omics data from both “female” and “male” samples become available. Hermaphroditic fish such as *Synbranchus marmoratus* and *Amphiprion perideraion* and invertebrates such as *Macrobrachium rosenbergii* and *Venus mercenaria* are candidates. In addition, we will add more omics data, such as sRNA-seq, BS-seq and ChIP-seq data. We expect that the resources in ASER will promote further studies to decode the molecular mechanisms of sex reversal.

## Data availability

ASER is publicly available at http://aser.ihb.ac.cn/.

## CRediT author statement

**Yangyang Li:** Software, Formal analysis, Data curation, Writing - original draft, Writing - review and editing. **Zonggui Chen:** Software, Formal analysis, Visualization, Writing - review and editing. **Hairong Liu:** Investigation, Data curation. Writing - review and editing. **Qiming Li:** Formal analysis. **Xing Lin:** Data curation. **Shuhui Ji:** Data curation. **Rui Li:** Data curation. **Shaopeng Li:** Data curation. **Weiliang Fan:** Visualization. **Haiping Zhao:** Data curation. **Zuoyan Zhu:** Writing - review & editing. **Wei Hu:** Conceptualization, Funding acquisition, Writing - review & editing. **Yu Zhou:** Conceptualization, Project administration, Funding acquisition, Methodology, Writing - review and editing. **Daji Luo:** Conceptualization, Project administration, Funding acquisition, Methodology, Writing - review and editing. All authors read and approved the final manuscript.

## Acknowledgements

This work was supported by grants from the National Natural Science Foundation of China (31922085, 31872191 to DL, 31922039 to YZ), the Strategic Priority Research Program of CAS (XDA24010108) to WH and DL, and Natural Science Foundation of of Hubei Province (2020CFA056 to DL and 2020CFA057 to YZ). Part of the computation of this work was done in the Supercomputing Center of Wuhan University and Hydrobiological Data Analysis Center.

## Supplementary material

**Figure S1.**
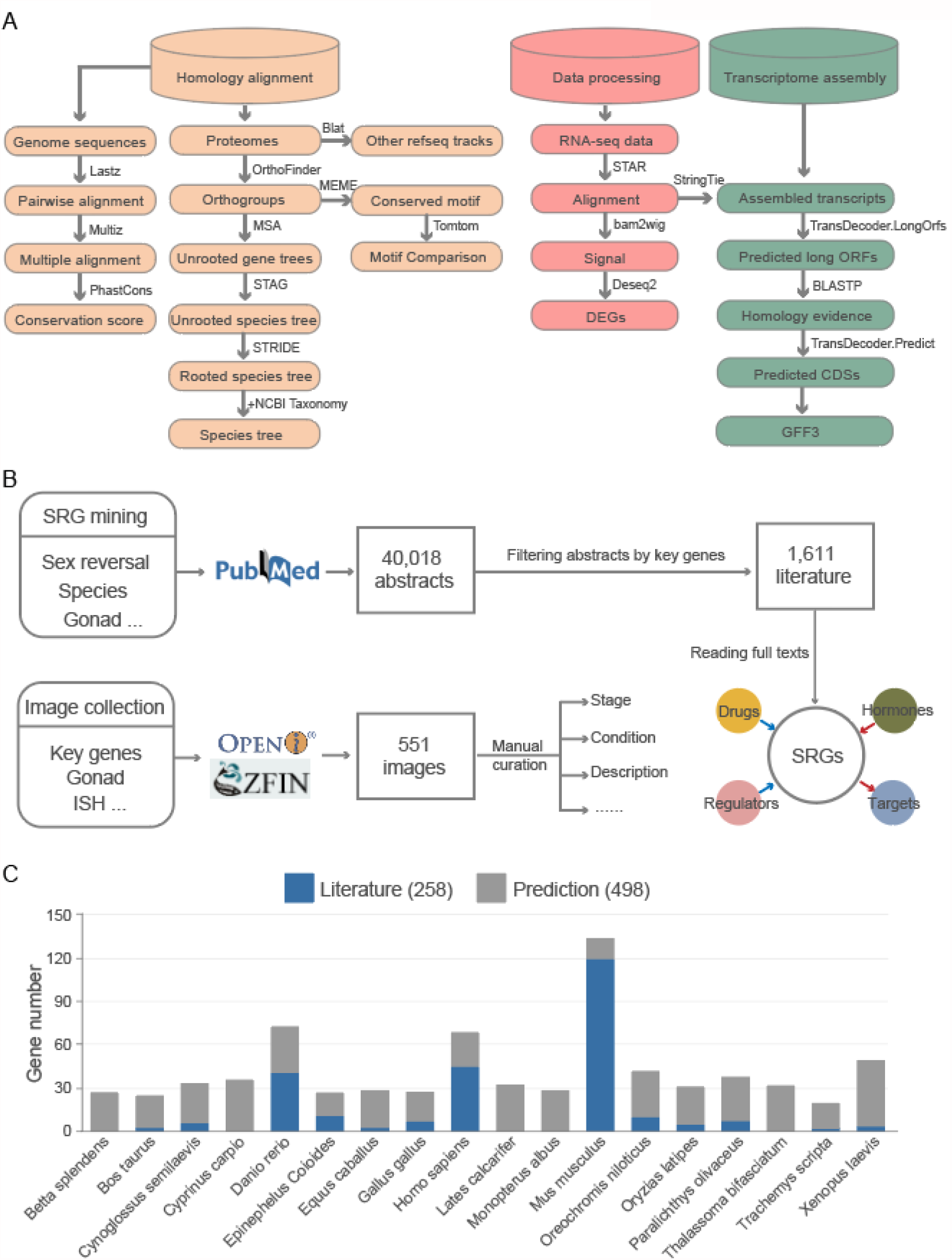
Data processing pipelines and statistics of SRGs. **A**. Workflows for building ASER, including homology alignment, RNA-seq data processing, and transcriptome assembly. **B**. Workflows for SRG mining, and image collection and annotation. **C**. Statistics of validated SRGs and predicted genes associated with sex reversal in the ASER database.

## Tables

**Table S1 RNA-seq data used in the ASER database**

**Table S2 Wordlist and blacklist in SRG mining**

**Table S3 Literature mining of SRGs for the ASER database**

